# Diverse flanking sequence types of *bla*_NDM_ in IncX3 and IncB/O/K/Z plasmids in *Escherichia coli* isolated from poultry

**DOI:** 10.1101/768622

**Authors:** Jia-Hang Zhu, Run-Mao Cai, Li-Juan Zhang, Xing-Run Zheng, Yue-Wei Lu, Qiu-Yun Zhao, Man-Xia Chang, Hong-Xia Jiang

## Abstract

We identified 33 *bla*_NDM_-harboring *Escherichia coli* in 470 poultry samples from Shandong and Guangdong provinces during 2016 to 2017 and included the subtypes *bla*_NDM-1_, *bla*_NDM-5_ and *bla*_NDM-9._ The *bla*_NDM_ gene possessed by all strains was plasmid-borne and ranged in size from 46 to 265 kb. The plasmids included the replicon types IncX3, IncY, IncB/O/K/Z and IncHI2 and most were transferable and stable in recipient host bacteria. Sequences flanking *bla*_NDM_ in IncX3 and IncB/O/K/Z plasmids were diverse and the mobile element IS*26* played an important role in the evolution of the IncB/O/K/Z *bla*_NDM_ plasmids.

---

Transmissible carbapenem-resistance and its dissemination in the *Enterobacteriaceae* have posed a major threat to global public health (1). In the existing 3 classes of carbapenemases, New Delhi metallo-beta-lactamases (NDM) encoded by *bla*_NDM_ is most abundant globally and its presence is increasing at an alarming rate in Asia (2, 3). The *bla*_NDM_ gene has been found primarily on plasmids with numerous replicon types including IncF, IncL/M, IncA/C, IncX, IncH, IncN, IncR, IncB/O and IncT (4, 5). In addition, this gene is often flanked by mobile genetic elements such as insertion sequences and transposons that are responsible for horizontal genetic exchange and therefore promote the acquisition and spread of resistance genes (4). The presence of carbapenemase-producing *Enterobacteriaceae* (CPE) in livestock animals is of particular concern because this may facilitate gene pool expansion from which pathogenic bacteria can pick up resistance genes, and consumers may subsequently be exposed through the food chain (6). The presence of *bla*_NDM_-harboring isolates is rapidly increasing among food animals in China and this is most likely related to the presence of *bla*_NDM_-harboring plasmids (7-13). We therefore examined *bla*_NDM_-harboring plasmids from poultry in Guangdong and Shandong provinces in China.

We collected 470 organ samples from five unrelated areas during 2016 in Shandong and 2017 in Guangdong (Table S1). *Escherichia coli* strains were isolated on MacConkey agar plates and were further identified by matrix-assisted laser desorption/ionization time of flight mass spectrometry (MALDI-TOF MS) and 16S rRNA gene sequencing (14). The *bla*_NDM_ gene was identified using the PCR method as previously described (15, 16). We found 33 *bla*_NDM_-harboring *E. coli* strains in our sample group and 9 were from Shandong and 24 were from Guangdong. Of the 33 strains, three *bla*_NDM_ subtypes were identified including *bla*_NDM-1_ (1/33), *bla*_NDM-5_ (24/33) and *bla*_NDM-9_ (8/33) (Fig. 1). Antimicrobial susceptibilities of these strains were determined using the agar dilution method according to CLSI guidelines. All 33 isolates were resistant to at least four classes of antibiotics including carbapenems (Table S2). Additional PCR screening for the presence of other resistance genes demonstrated that all *bla*_NDM_-harboring *E. coli* co-harbored other resistance genes and 10 of which co-harbored polymyxin resistance gene *mcr-1* (Table S3 and Fig 1). Pulsed-field gel electrophoresis (PFGE) analysis identified the presence of 19 clusters at 85% identity (17) (Fig 1).

**Figure 1.**
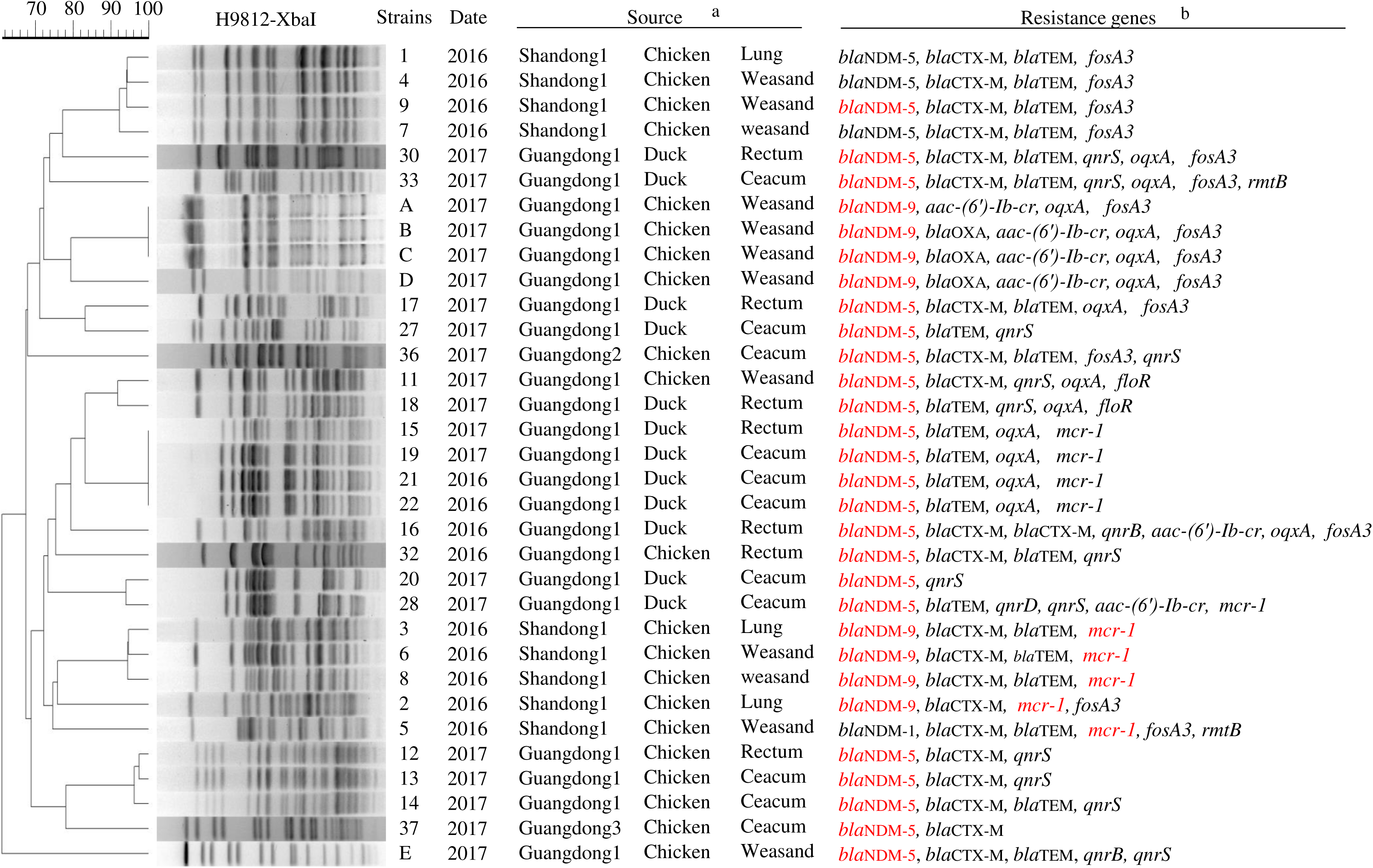
PFGE patterns and resistance genes of 33 *bla*_NDM_-harboring *E. coli* strains. ^a^ Shandong1, Guangdong1, Guangdong2 and Guangdong3 represent four different areas in Shandong and Guangdong provinces. ^b^ Genes shown in red color represent that it could be transferred through conjugation.

Conjugation assays were performed with the 33 *bla*_NDM_ strains as previously described using *E. coli* J53 as the recipient (18). Transconjugants harboring *bla*_NDM_ and *mcr-1* were selected on MacConkey agar / sodium azide (150 µg/mL) plates that also contained either meropenem (0.3 µg/mL) or colistin (0.5 µg/mL), respectively. All transconjugants were confirmed by PCR and Enterobacterial repetitive intergenic consensus (ERIC) PCR assays (19). The *bla*_NDM_ gene from 29/33 strains was successfully transferred, as was *mcr-1* in 5/10 of the *mcr-1*-harboring strains (Fig. 1 and Table 1). The plasmids carrying *bla*_NDM_ were characterized by PCR-based replicon typing (PBRT) (20, 21), S1 nuclease PFGE and Southern hybridization (18, 22). The *bla*_NDM_ genes were all located on plasmids of different replicon types that included IncB/O, IncY, IncX3 and IncHI2 with sizes from 46 to 265 kb (Table 1).

**Table 1.**
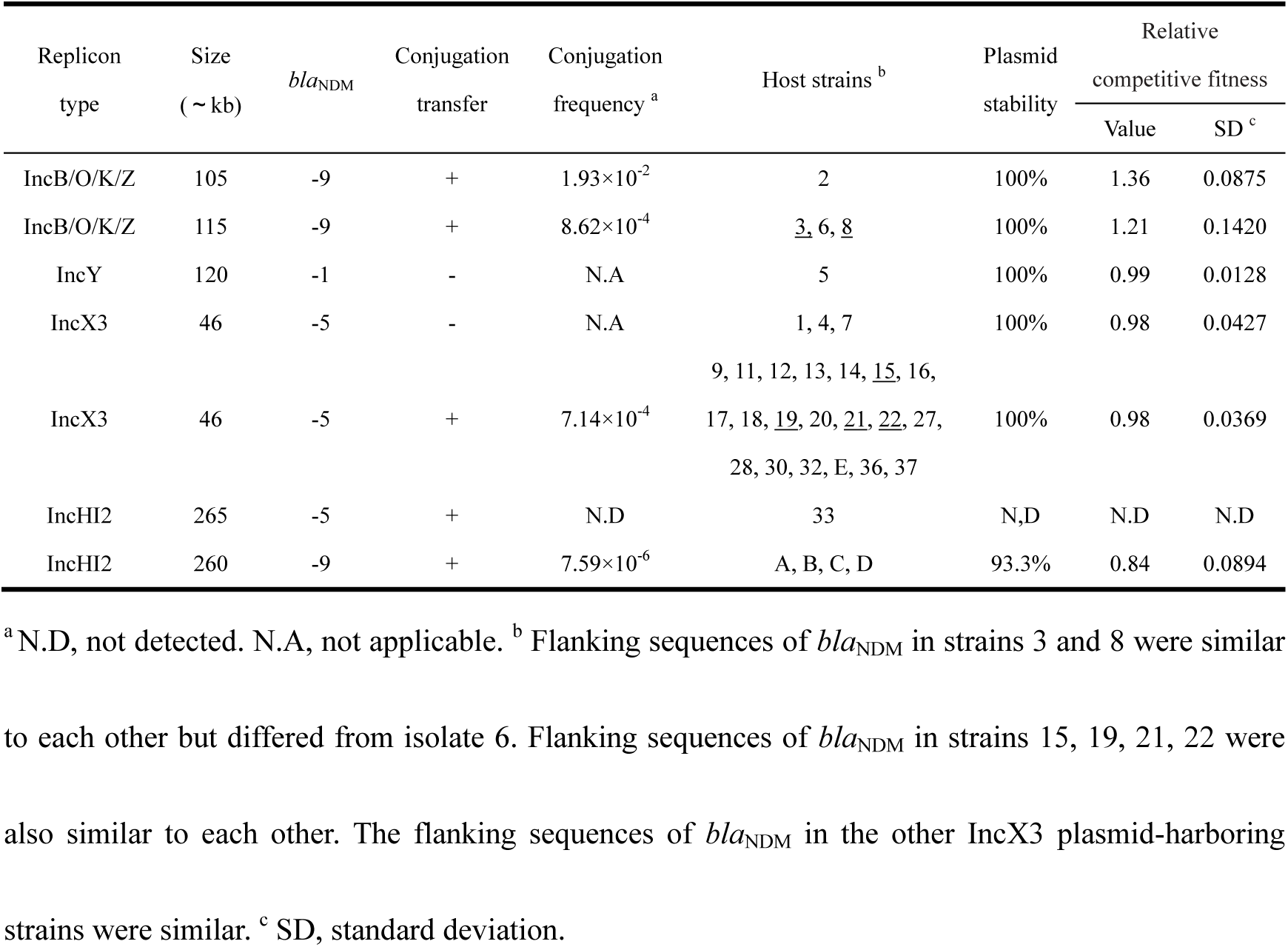
Characteristics of *bla*_NDM_-harboring plasmids among *E. coli* isolates.

The fitness of different *bla*_NDM_-harboring plasmids was tested *via* stability and competition experiments as previously described (23, 24). Plasmid DNA from transconjugants were extracted using plasmid minikits (Qiagen, Hilden, Germany) and then electroporated into *E. coli* DH5α except for the plasmid from strain 33 that failed. Each transformant was confirmed to contain a single plasmid by S1 nuclease PFGE. Transconjugants harboring *bla*_NDM_ were selected on Luria-Bertani agar plates containing cefotaxime (1 µg/mL) and incubated 8 days for stability assays and 12 days for competition assays. Except for the replicon HI2 type plasmids, the remainders were stable and persistent in their host strains without reduced fitness (Table 1), these are conditions conducive to the dissemination of *bla*_NDM_.

We sequenced the *bla*_NDM_-harboring plasmids of the replicon IncB/O type (105 and 115 kb) and the IncX3 type using the Illumina Hiseq platform (Majorbio, Shanghai) and assembled data using SOAP denovo. Gaps were closed through PCR and Sanger sequencing. Plasmid sequences were analyzed using BLAST (http://blast.ncbi.nlm.nih.gov/Blast.cgi) and plasmid replicon types were analyzed using the PlasmidFinder tool (https://cge.cbs.dtu.dk/services/PlasmidFinder/). PCR primers were designed to detect the *bla*_NDM_ genetic environment of similar plasmids (Tables S4 and S5).

Two types of sequences flanking *bla*_NDM_ were found in the IncX3 plasmids that were assigned the names pNDM-T16 and pNDM-T21. Plasmid pNDM-T16 was 46161 bp and almost 100% identical to plasmid pNDM5_IncX3 (KU761328) with only 6 single-base changes, in which the IS*Aba125* sequence adjacent to *bla*_NDM-5_ was interrupted by IS*5* and split into two fragments. The sequence of pNDM-T21 has not been reported previously and downstream of IS*5* was a truncated IS*3000* transposase gene that most likely was the product of intramolecular recombination (Figure S2). This discovery gives further evidence for the diversity of the sequence types for IncX3 *bla*_NDM_-harboring plasmids as reported previously (25-27).

The IncB/O plasmids were identified as IncB/O/K/Z plasmids using the PlasmidFinder tool. Three kinds of sequences flanking *bla*_NDM_ were found and the plasmids were assigned the names pNDM-T2, pNDM-T3 and pNDM-T6. The major genetic organization surrounding *bla*_NDM-9_ in these plasmids was similar to that in the plasmids pHNTH02-1 (MG196294) and p5CRE51-NDM-9 (CP021177). All 5 plasmids possessed IS*26*-*intI1*-*dfrA12*-*qacEΔ1*-*sul1*-IS*CR1*-(ΔIS*1294*)-Δ*dct*-*tat*-*trpF*-*ble*_MBL_-*bla*_NDM_-ΔIS*Aba125*-IS*26* and this IS*26* composite transposon was most likely responsible for *bla*_NDM_ transfer (28) (Fig. 2).

**Figure 2.**
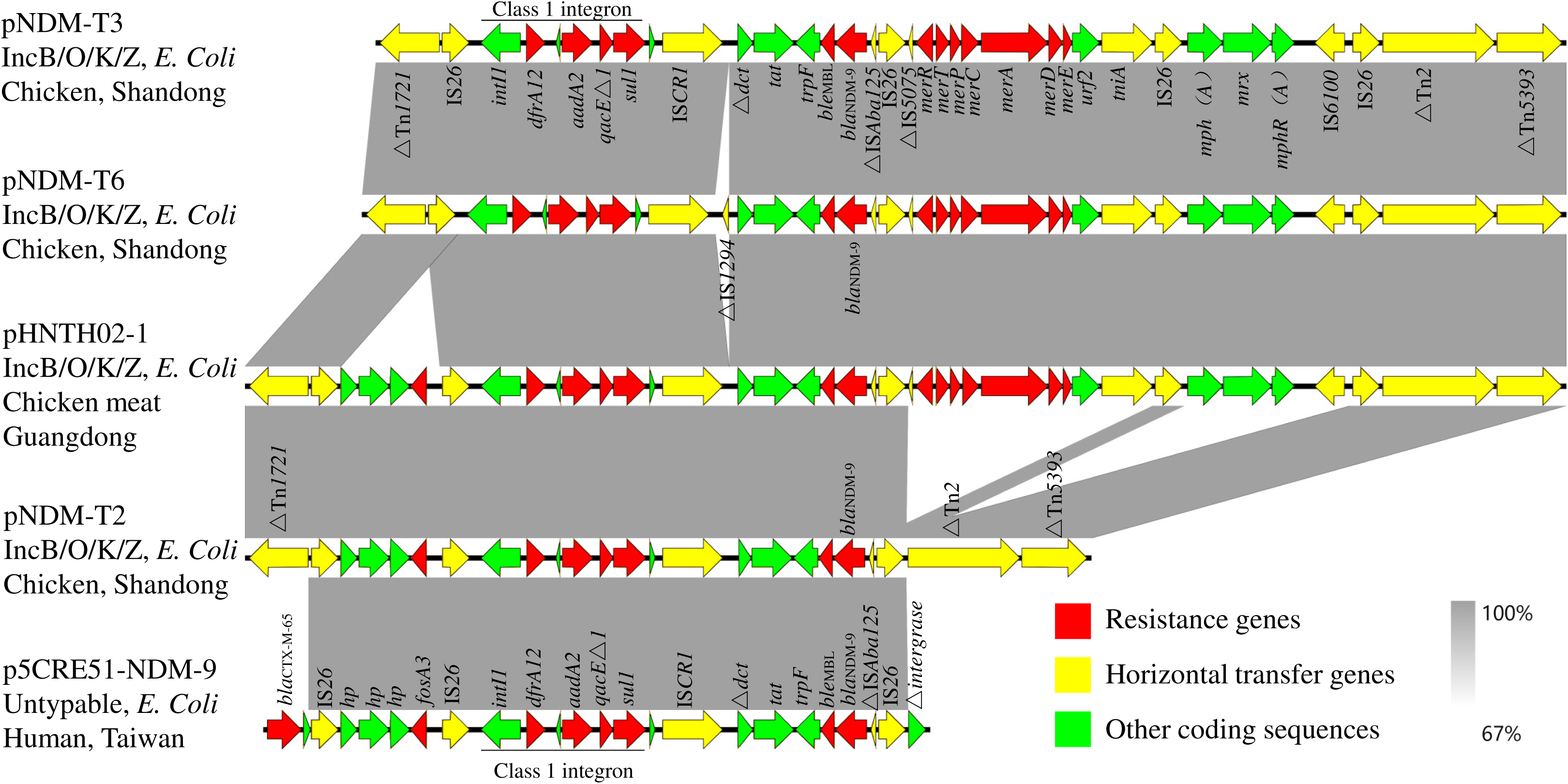
Comparison of the *bla*_NDM_ genetic environment in pNDMT-2 and pNDMT-6 with other plasmids.

The sequences surrounding the *bla*_NDM-9_ genes were diverse and was most likely the result of the action of the IS*26* mobile element. We therefore designed specific PCR primers to detect circular intermediates (circle A-D) and other possible structures (structure A-G) mediated by IS*26* in three IncB/O/K/Z plasmids (Table S6 and Fig3). The PCR products were confirmed by Sanger sequencing and we found only one cyclic intermediate formed by IS*26*-*mph(A)*-*mrx*-*mphR(A)*-IS*6100*-IS*26*. Interestingly, the A-G structures were all detected although no direct repeats were found adjacent to the IS*26* insertions. These results demonstrated there were various plasmids possessed different genetic organizations surrounding *bla*_NDM-9_ in the host bacteria, and all were derived from a larger plasmid *via* IS*26* transposition. Remarkably, even though the circular intermediates containing *bla*_NDM_ were not detected in these plasmids, IS*26*-*intI1*-*dfrA12*-*qacEΔ1*-*sul1*-IS*CR1*-Δ*dct*-*tat*-*trpF*-*ble*_MBL_-*bla*_NDM_-ΔIS*Aba125*-IS*26* has also been found in IncA/C2 (pNDM-2248, CP021177), IncHI2-N (pC629, CP015725) and IncFII(Y)-like (pKPGJ-1a, CP017283.1) plasmids that originated from *E. coli, Salmonella* Indiana, and *Klebsiella variicola*, respectively (28, 29). This suggests the possible transfer of *bla*_NDM_ between different plasmids mediated through IS*26*-mediated composite transposon.

**Figure 3.**
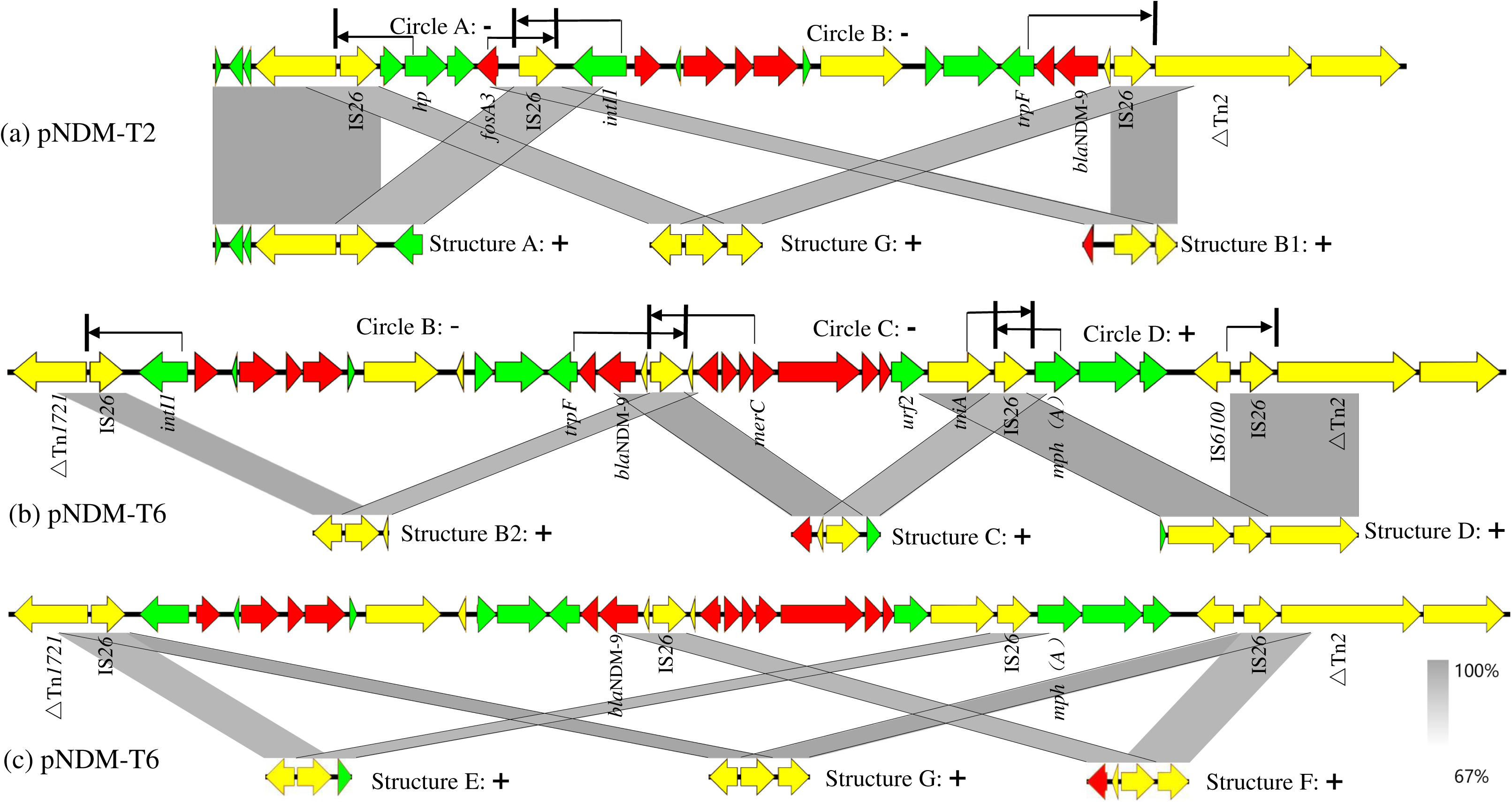
Circular intermediates and possible structures present in the IncB/O/K/Z plasmids found in this study.

In conclusion, this study demonstrated that the dissemination of *bla*_NDM_ among poultry is closely related to diverse plasmids, especially the IncX3 and IncB/O/K/Z conjugative plasmids. The sequences flanking *bla*_NDM_ in IncB/O/K/Z as well as in IncX3 plasmids, tended to be diverse and was most likely the result of intramolecular recombination and frequent activity of mobile elements such as IS*26*. These events resulted in new types of *bla*_NDM_-harboring plasmids and are related to the process of evolution for the *bla*_NDM_-harboring plasmids over time. Given the continued spread of *bla*_NDM_ gene and the significant role that mobile elements and cyclic intermediates played in the evolution and dissemination of resistance plasmids (2, 30), further studies concerning the molecular mechanisms behind the dissemination of *bla*_NDM_ are urgently needed.

## Accession numbers

The complete nucleotide sequences of the three *bla*_NDM_-carrying plasmids characterized in this study were submitted to GenBank under accession numbers MN335919 (pNDM-T2), MN335921 (pNDM-T6), MN335922 (pNDM-T16), and the annotated sequences of the *bla*_NDM_ genetic environment identified from strain 3 and strain 21 have been submitted to GenBank under accession number MN335920 (pNDM-T3), MN307121 (pNDM-T21), respectively.

## SUPPLEMENTAL MATERIAL

Supplemental file 1, PDF file, ……MB.

## ACKNOWLEDGMENTS

This work was supported in part by the National Natural Science Foundation of China (31972734) and National key research program of China (grant 2016YFD0501300).

---

**Figure S1.**
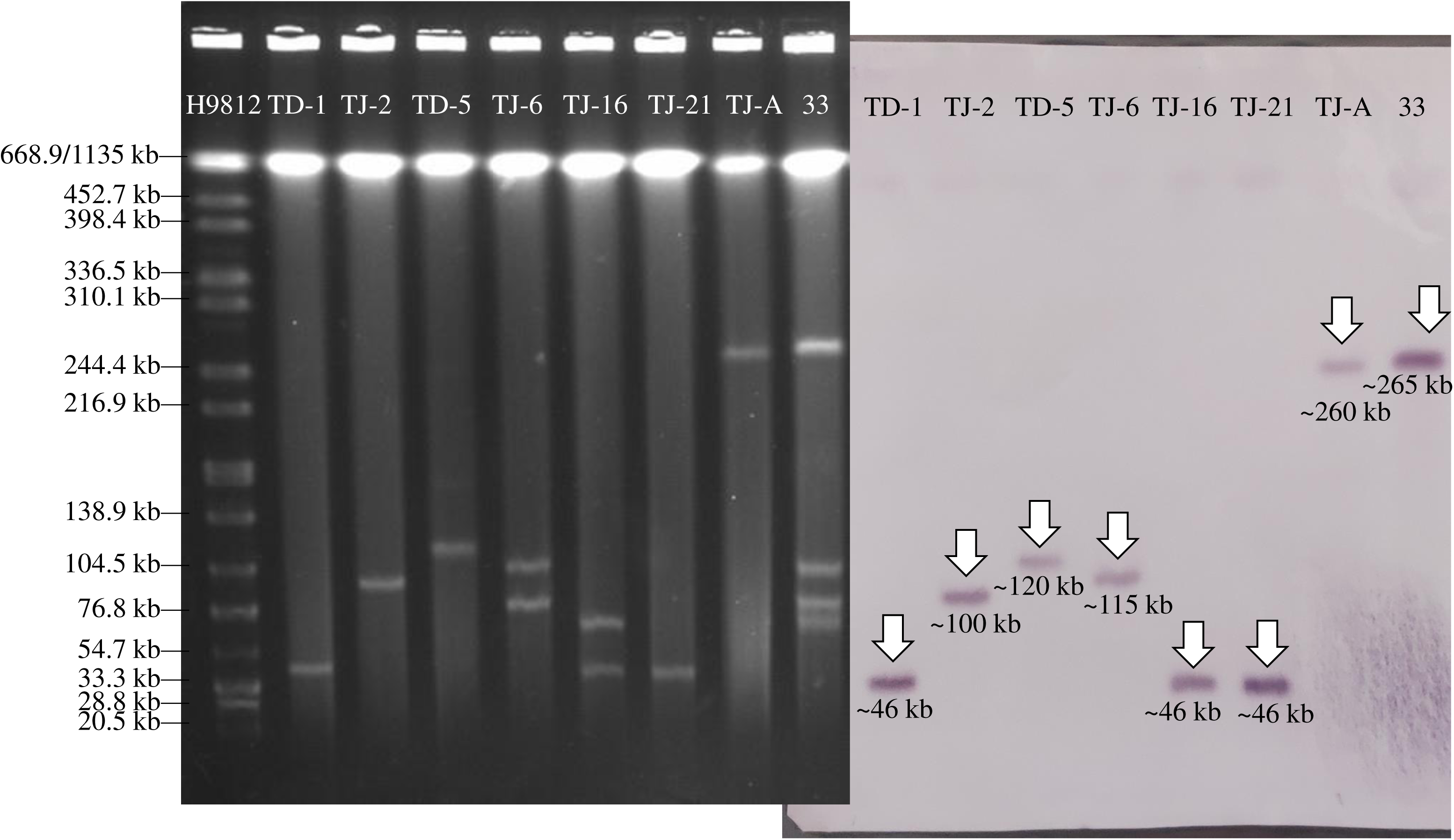

**Figure S2.**
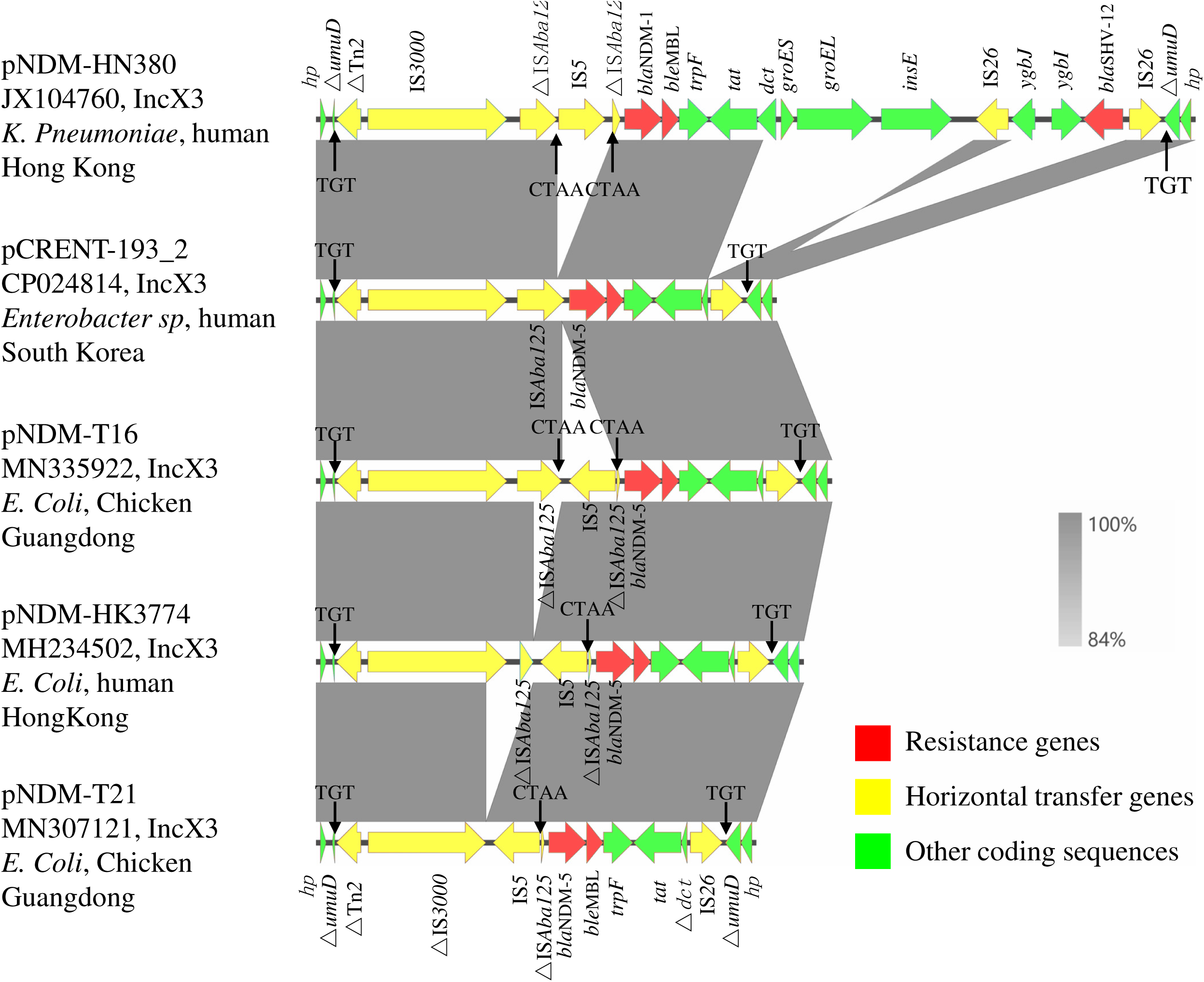

**Figure S3.**
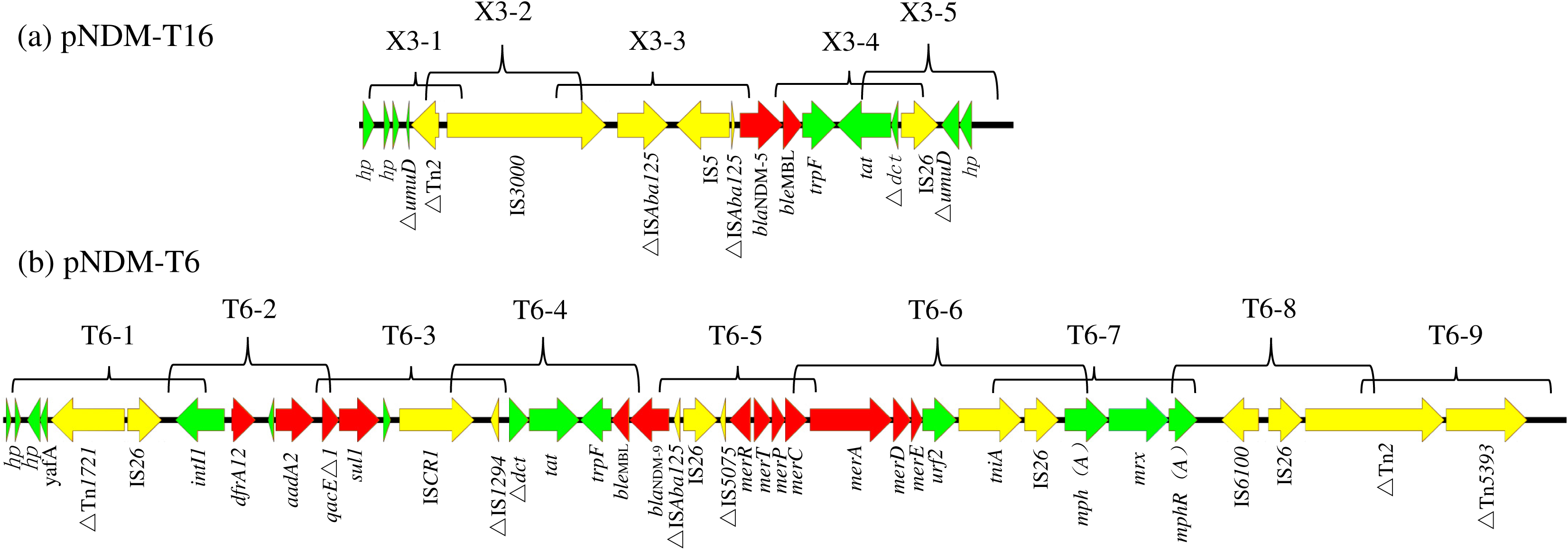

